# iProMix: A decomposition model for studying the function of ACE2 based on bulk proteogenomic data for coronavirus pathogenesis

**DOI:** 10.1101/2021.05.07.441534

**Authors:** Xiaoyu Song, Jiayi Ji, Pei Wang

## Abstract

Both SARS-CoV and SARS-CoV-2 use ACE2 receptors to enter epithelial cells in lung and many other tissues to cause human diseases. Genes and pathways that regulate ACE2 may facilitate/inhibit viral entry and replication, and genes and pathways that are controlled by ACE2 may be perturbed during infection, both affecting disease severity and outcomes. It is critical to understand how genes and pathways are associated with ACE2 in epithelial cells by leveraging proteomic data, but an accurate large-scale proteomic profiling at cellular resolution is not feasible at current stage. Therefore, we propose iProMix, a novel framework that decomposes bulk tissue proteomic data to identify epithelial cell component specific associations between ACE2 and other proteins. Unlike existing decomposition based association analyses, iProMix allows both predictors and outcomes to be impacted by cell type composition of the tissue and accounts for the impacts of decomposition variations and errors on hypothesis tests. It also builds in the functions to improve cell type estimation if estimates from existing literature are unsatisfactory. Simulations demonstrated that iProMix has well-controlled false discovery rate and large power in non-asymptotic settings with both correctly and mis-specified cell-type composition. We applied iProMix to the 110 adjacent normal tissue samples of patients with lung adenocarcinoma from Clinical Proteomic Tumor Analysis Consortium, and identified that interferon *α* and *γ* pathways were most significantly associated with ACE2 protein abundances in epithelial cells. Interestingly, the associations were sex-specific that the positive associations were only observed in men, while in women the associations were negative.

## 1 Introduction

In the past two decades, three types of coronaviruses emerged to cause serious and widespread human illness and death, including SARS-CoV (2002), MERS-CoV (2004) and SARS-CoV-2 (2019). While MERS-CoV used DPP4 receptor to enter human cells, both SARS-CoV and SARS-CoV-2 used the same ACE2 receptor for cell entry [1]. The abundance of ACE2 receptor on the cell surface has been identified as the limiting factor of viral attachment, fusion and entry [2], and thus affects the viral replication rate and disease severity [3]. The upstream genes and pathways that regulate the abundance of ACE2 receptor may facilitate or inhibit the virus infection to host cells, and the downstream genes and pathways that are regulated by ACE2 receptors may be perturbed when the receptor is bounded with the virus and cannot perform its regulatory functions. Therefore, identifying genes and pathways associated with ACE2 receptor paves the way for understanding the viral pathogenesis and the host defense mechanisms to prevent and treat the severe illness. Tools that identify these associations are not only for useful understanding current COVID-19 pandemic but also prepare us for new types of coronavirus potentially emerging in the future.

The ACE2 was primarily expressed in epithelial cells [1, 2, 4]. A recent study that leveraged the single cell RNA sequencing (scRNAseq) data of airway epithelial cells identified the interferon (IFN)*α* and INF*γ* pathways may stimulate ACE2 expression levels [4]. This result brought concerns over the complex and contradictory role of IFNs and the general immune system in combating coronaviruses: on one side it induces interferon-stimulated genes to promote host antiviral defense, but on the other side it may also induce ACE2 which makes the virus easier to enter the cells and replicate. Validating the result is critical to generate insights for clinical prescription of IFN-based therapies and development of novel therapeutics. As a result, a validation experiment was performed by perturbing IFN in cell lines, which although resulted in differential expression of ACE2, the effects on ACE2 protein abundances were not identified [4]. Proteins are critical molecules that carry out almost every single cellular function. They are poorly represented by gene expressions as proteins employee complex, multi-level post translational modification that are not reflected at the mRNA level. Studies that quantified both mRNA gene expressions and proteomic abundances identified a median correlation of <0.5 in lung [5] and many other tissues [6, 7]. Therefore, identifying and validating genes and pathways at protein level that are associated with ACE2 is crucial for further understanding the roles of this gene in coronavirus infection.

Unlike scRNAseq, the proteomic profiling technology at the cellular resolution is not mature enough for large-scale studies. Analyses at tissue level have been confounded by other cell types [5]. On the other hand, statistical methods for decomposing bulk -omic data have been intensively studied these years [8, 9, 10, 11, 12, 13, 14, 15, 16, 17, 18, 19, 20, 21, 22, 23, 24] and shed light on developing statistical methods for studying cell-type specific associations with ACE2. The existing decomposition methods can be broadly categorized into two groups based on their goals. The first group focuses on estimation, such as estimating the cell-type composition [14, 17, 18] and the cell-type specific transcriptomic levels averaged across samples or for each individual sample [25, 20]. The second group focuses on associations by leveraging pre-estimated cell-type composition for cell-type controlled or specific effects. For example, cell-type composition estimates are often treated as covariates in simple association analysis for “unconfounded” effects [6, 7]. Cell-type specific associations are also considered by modeling interactive effects between the estimated cell-type composition and covariates [26] or by considering hierarchical mixture models to decompose the observed tissue-level traits into their values in different cell types for their cell-type specific associations [27]. There are a few difficulties for applying existing methods to our study to identify genes and pathways associated with ACE2 abundances in epithelial cells: First, the protein abundance (or gene expression) levels of both ACE2 and other genes are impacted by the cell type composition. Existing methods decompose the outcomes and regress them on the known predictors. In our setting both predictor (e.g. ACE2) and outcome (e.g. other gene) need to be decomposed. Secondly, existing cell-type specific association analyses assumed the cell-type composition as known and did not account for its estimation biases and variations in hypothesis tests. However, the cell-type compositions are estimated and how the estimation would affect the test statistics is not well studied and accounted, and thus the results may suffer from uncontrolled false positives and reduced study power. Lastly, the estimation for cell-type composition is an ongoing effort that is far from achieving perfection. For example, most well-established methods estimate cell type proportions for a limited number of cell types and epithelial cells are not among the most popular cell types. The xCell [17] is a method that considers 64 cell types including epithelial cells, but it provides only an estimation for relative abundances instead of proportions of a cell type. In other words, it describes how the abundance of epithelial cells in one sample relative to other samples, instead of proportion that tells how the abundance of epithelial cells compared to other cell types within the same sample.

In this study, we propose a new statistical method *iProMix* to analyze the association between ACE2 and other proteins in epithelial cells by leveraging proteogenomic data in bulk tissue. iProMix is building upon a mixture model which treats the observed bulk profiles of both ACE2 and other proteins as a weighted average from epithelial and non-epithelial cell types. It models the strength of cell-specific associations via their conditional joint distributions, and accounts for the uncertainty of decomposition in hypothesis testing. It leverages existing estimation on cell-type composition but builds in an option to allow an updated estimation from the data. We applied iProMix to adjacent normal tissue of lung adenocarcinoma (LUAD) from Clinical Proteomic Tumor Analysis Consortium (CPTAC), and confirmed the role of proteins in IFN*α* and IFN*γ* pathways on the ACE2 expression and protein abundance levels in epithelial cells. These associations were not observed by looking at the tissue level data without decomposition. Strikingly, we observed sex-specific effects that were not observed in previous studies.

The article is structured as follows: Section 2 describes the iProMix model that identifies cell specific associations between ACE2 and other proteins in epithelial cells with bulk tissue data. Section 3 presents various simulation results for the performance of our approach in comparison with traditional tissue-level models for error control and study power in correctly and mis-specified models. Section 4 presents the application to CPTAC LUAD data with a focus on IFN*α* and IFN*γ* pathways.

## 2 Method

### 2.1 Model

In this section, we introduce the proposed iProMix, an *i*ntegrative *Pro*teogenomic model for bulk tissues that are a *Mix*ture of different cell types. Suppose a data set has proteomic profiling of *N* tissues. In tissue *i* (*i =* 1, …, *N*), we let *x*_*i*_ be the ACE2 protein abundance (or expression level), *y*_*i*_ be the protein abundance of a different gene, and ***z***_*i*_ be a set of covariates such as age. iProMix decomposes the observed protein abundance (or gene expression) levels into their unobserved levels in epithelial-cell component (*x*_*ui*_, *y*_*ui*_)and non-epithelial cell components (*x*_*vi*_, *y*_*vi*_). Similar to many models [24, 27], we assume that the observed tissue-level value is a linear combination of their values in the two major cell types. We have

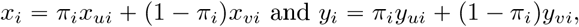

with *π*_*i*_ denotes the proportion of epithelial cells in the tissue.

As both *x*_*ui*_ and *y*_*ui*_ can not be directly observed, regression-based framework cannot work for identifying their associations. We propose to leverage a joint modeling idea to identify their conditional correlation given ***z***_*i*_. In details, we model their dependence structures via multivariate Gaussian distribution as follows: ***u***_*i*_ *=*(*x*_*ui*_, *y*_*ui*_)^*T*^ ∼ *N* (***µ***_*ui*_, Σ_*u*_) and ***v***_*i*_ *=* (*x*_*vi*_, *y*_*vi*_)^*T*^ ∼ *N* (***µ***_*vi*_, Σ_*v*_), where the mean functions depend on covariates ***z***_*i*_ as such 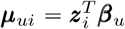 and 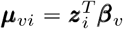, and the variance-covariance functions 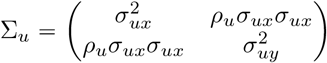 and 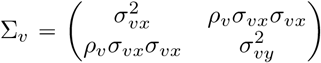 quantify their dependence structure in the epithelial- and non-epithelial components, respectively. Collectively, 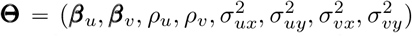 and ***π*** {*π*_1_,…, *π*_*n*_} are the model parameters. The hypothesis test for the association between ACE2 and protein abundance given ***z***_*i*_in epithelial cell type becomes to test *H*_0_ : *ρ*_*u*_ *=* 0 v.s. *H*_*a*_ : *ρ*_*u*_ ≠ 0.

### 2.2 Estimation of *π*

We propose to estimate epithelial cell proportion ***π*** separately from **Θ**, as ***π*** is a feature of the tissue that remains the same regardless of the genes, -omic data types and inclusion of other tissues. As a result, we can use -omic data from other technologies (e.g. RNAseq) that are more established and have more prior information than proteomic profiling to improve the estimation accuracy. Among many methods that are available for estimating cell-type composition, most are based on cell-type signature genes, which are not specific to the tissue types and sample conditions (e.g. healthy vs. diseased). Therefore, we employ a hybrid approach to both take advantages of existing knowledge and extract additional signals from the data for cell-type composition estimation. We assume there exists a prior estimate of epithelial cell proportion *h*_*i*_ for tissue *i* from independent sources, and use *Beta* distribution to link it to the true *π*_*i*_ as done in previous studies [24]. Specifically, we assume *h*_*i*_ ∼ *Beta* (*α*_*i*_, *β*_*i*_), where *α*_*i*_ *= π*_*i*_*δ* and *β*_*i*_ *=* (1 − *π*_*i*_)*δ* for some positive parameter *δ*. By assumption, *E* (*h*_*i*_)*= π*_*i*_ and *var* (*h*_*i*_)*= π*_*i*_ (1 − *π*_*i*_)*/*(*δ* +1) that *h*_*i*_ is an unbiased estimator of the true level of *π*_*i*_. Suppose we use *G* individual genes collected on a -omic data type of *M* tissue samples to estimate 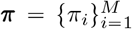 and *δ*. These *M* sample should include *N* samples used in downstream analysis but can also include additional samples (e.g. sample with with RNAseq but not proteomic data) to improve estimation accuracy. We let 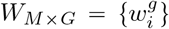 be the observed profiling of *M* samples and *G* genes in the tissue, 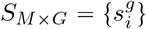 be the unobserved levels in epithelial cells and 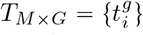 be the unobserved levels in non-epithelial cells. Under the assumption that the observed tissue level is as a linear combination of levels in the epithelial- and non-epithelial-cells, we have

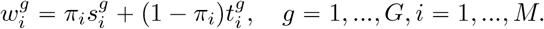

As the primary goal is to estimate ***π*** instead of capturing the interrelationship among the genes, we consider the marginal distribution of the genes ignoring their correlations to allow a computationally efficient estimation. We have

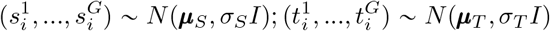

with *I* being the *G* × *G* dimensional identity matrix. Assuming independence between {*h*_*i*_} and 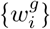, the estimation process is the solution of the following maximization problem:

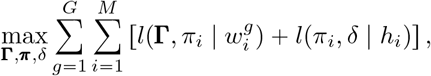

where **Γ** = (***µ***_*S*_, ***µ***_*T*_, *σ*_*S*_, *σ*_*T*_), *δ* and ***π*** are the model parameters. The 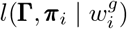 and *l* (***π***_*i*_, *δ* |*h*_*i*_)are the log likelihood of the observed gene expression profile and epithelial cell proportion estimate, respectively. Given this likelihood function, we carry out a generalized Expectation-Maximization (EM) algorithm to estimate ***π*** and *δ* which can be summarized in the following two steps:

1. E-step: Given the current estimates of the model parameters, i.e. ***π***^(*t*)^, *δ*^(*t*)^, **Γ**^(*t*)^, we calculate 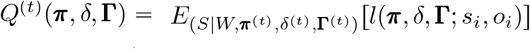.
2. M-step: We sample each parameter ***π***, *δ*, **Γ** conditioning to other parameters via the Expectation Conditional Maximization (ECM) algorithm [28] to calculate 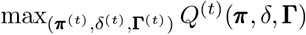.

As the estimated parameters depend on prior, input genes and samples, we propose bootstrap procedure to use different priors, genes and samples for estimation, and aggregate the multiple estimates for a robust estimation. We propose to calculate a tail truncated mean for aggregation to minimize the impact of outliers. It is worthy noticing that as *E* (*h*_*i*_)*= π*_*i*_, the averaged value of final estimates 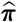 depends on the mean of prior, and thus the downstream analysis should be robust against a mis-specification of the averaged prior.

### 2.3 Estimation of Θ

We propose a likelihood based method to estimate the parameter **Θ** in iProMix assuming 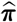 is known. The complete data log-likelihood function, *l*, of the observed data ***o***_*i*_ = (*x*_*i*_, *y*_*i*_) and unobserved data ***u***_*i*_ and 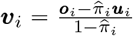 with respect to the parameter **Θ** can be expressed as follows:

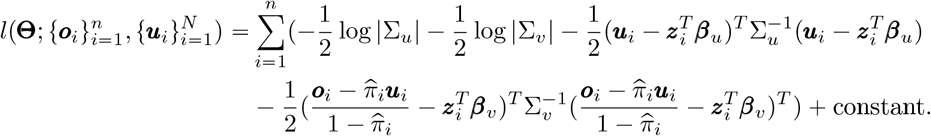

The identifiability of the parameter **Θ** is demonstrated in Supplementary Materials. If ***u***_*i*_ and ***v***_*i*_ are observed, maximum likelihood estimates of the parameters **Θ** could be directly obtained. The latent (unobserved) nature of ***u***_*i*_ and ***v***_*i*_ requires the adoption of the EM algorithm. Specifically, the EM algorithm summarizes into the following steps:

- E-step: Given the current estimates of the model parameters, i.e. **Θ**^(*t*)^, we calculate 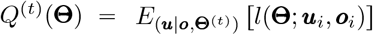.
- M-step: We find **Θ**^(*t* + 1)^ which is the solution to the following max_**Θ**_ *Q*^(*t*)^(**Θ**).

In the EM algorithm, a good initialization can lead to faster convergence than random starts. We assumed *ρ*_*u*_ and *ρ*_*v*_ to be zero and used the the mean and variance from regressing ***o***_*i*_ on ***z***_*i*_ as the initial values.

### 2.4 Two hypothesis test strategies

Building upon parameter estimation, the goal of iProMix is to identify proteins associated with ACE2 in epithelial cells. If the cell-type level data is observed, the hypothesis test of *ρ*_*u*_ 0 can be solved by likelihood ratio test (LRT), which approximately follows a 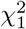 distribution under the null hypothesis. iProMix, however, considers a decomposition model that includes unobserved data and leverages quantities such as 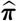 that are estimated with errors and variations. Therefore, in addition to iProMix (*χ*^2^), we consider two empirical distribution based strategies, iProMix (Permutation) and iProMix (Knockoff), to leverage the parallel nature of the genes in -omics data types to identify individual genes with controlled false discovery rate (FDR). This idea borrows information across genes to assist inference about each gene individually, which has been extensively employed in -omics studies, such as in FDR [29] and limma [30].

Suppose we consider a total of *P* other proteins 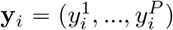 for their association with ACE2 *x*_*i*_. In both approaches, we permute the label ACE2 to generate 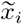 that preserves the distribution of ACE2 but breaks its correlation with **y**_*i*_’s. The 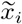 copy behaves in the same way as the original variables but, unlike them, is known to be null. How we handle 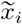 in the two strategies are, however, different. The results for iProMix (Permutation) and iProMix (Knockoff) are non-asymptotic.

#### iProMix (Permutation)

We calculate the LRT statistics in original and permuted separately, and compare their values for FDR controlled discoveries. In original data for protein *p*, we estimate the full log likelihood function of 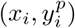 as 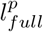,and the reduced log likelihood function 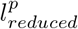 by forcing 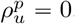. The corresponding LRT statistic for 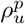 is defined as 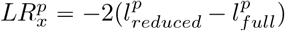. Similarly, in permuted data, we model 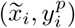 and get its LRT statistic 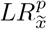 Across all *P* proteins, we consider a pre-specified LRT cut-off value *c* and a protein was counted if its LRT statistics is > *c*. We calculated empirical FDR (eFDR) as

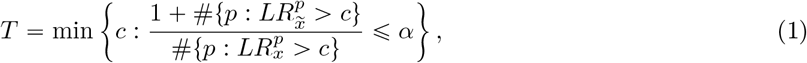

where *α* is a pre-determined nominal FDR level. By the definition of false discovery proportion (FDP), we have 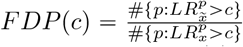, and incrementing the number of numerator by one the slightly more conservative procedure in Equation (1) controls the FDR.

#### iProMix (Knockoff)

The null distribution for 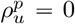 includes two cases: (1) both 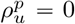 and 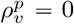, and (2) 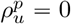 but 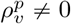. While iProMix (Permutation) provides a negative control for ACE2 and protein association in epithelial cells, it also breaks the ACE2 and protein association in non-epithelial cells, and thus its null distribution does not include case (2). To overcome this issue, we consider an alternative strategy iProMix (knockoff) to model the joint distribution of 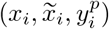 in iProMix framework. Assume the observed data can be decomposed into the epithelial component 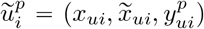 and non-epithelial component 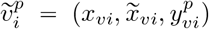, and their cell-type specific dependence can be modeled by multivariate Gaussian distribution similar as described above. Then 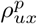 and 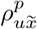 denote the 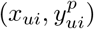 and 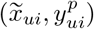 correlation conditional on other variables in the model, respectively. A reduced model for 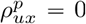 gives a log-likelihood function 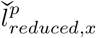 and a reduced model for 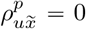 gives a log-likelihood function 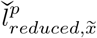. Then, we define the iProMix (Knockoff) test statistic as the difference of two LRTs that

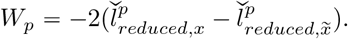

When protein *p* is under the null, 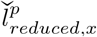 is close to 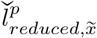 and *W*_*p*_ is close to 0. *W*_*p*_ > 0 indicates that the association of protein *p* with ACE2 is more important than with its knockoff copy. Hence a large positive value of *W*_*p*_ is an indication that this protein is a genuine signal and associated with ACE2 in the model. Finally, we leverage the empirical distribution of from all *P* proteins to control the FDR. For a pre-specified cut-off value *c* >0, protein *p* was considered positive if its *W*_*p*_ is >*c*. The FDR controlled cutoff of *W*_*p*_ is defined as

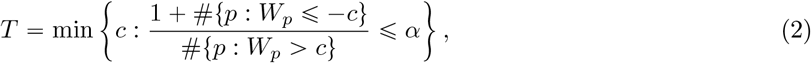

where *α* is the pre-determined nominal FDR level. This equation employs the property of *W*_*p*_ that it is symmetric under the null, and for any fixed threshold *c* >0, we have # {*p* : *W*_*p*_ ⩽ −*c*}= #{null *p* : *W*_*p*_ ⩽ − *c*} = #{null *p* : *W*_*p*_ ⩽*c*}. By incrementing the negatives by one, the slightly more conservative procedure controls the FDR.

### 2.5 Pathway enrichment analysis

One can perform pathway enrichment analysis with significant genes, but this analysis is sensitive to the power and error control of gene identification. Alternatively, we propose to compare the relative rank of *LRT* statistics for all proteins, which is robust against to the distribution assumptions of LRTs that are perturbed by decomposition. For protein *p*, we define *Q*^*p*^ as the rank of its signed LRT score across all proteins as

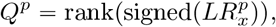

where the sign comes from the estimated correlation 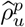 of the data, and the 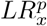 is the likelihood ratio test statistic of contrasting full and reduced models of 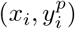. For a pathway, we let ***Q***^*in*^ indicate the scores for all proteins inside of the pathway, and ***Q***^*out*^ indicate scores for all proteins outside of the pathway. Then, we compared ***Q***^*in*^ versus ***Q***^*out*^ such as using non-parametric Wilcoxon rank score test to identify ACE2 enriched pathways.

Suppose a pathway is not associated with ACE2, then ***Q***^*in*^ and ***Q***^*out*^ present similar distributions. If ***Q***^*in*^ is significantly greater than ***Q***^*out*^, it indicates this pathway is positively associated with ACE2. Similarly, if ***Q***^*in*^ is significantly smaller than ***Q***^*out*^, it indicates this pathway is negatively associated with ACE2. We considered 50 pathways known from Hallmark database [31], including INF*α* and IFN*γ*, and considered Benjamini–Yekutieli (BY) procedure for FDR control due to highly overlapped genes across pathways.

## 3 Simulation results

### 3.1 Finite sample performance under correctly specified *π*

In this section, we evaluated the FDR and power of iProMix methods for identifying cell-specific associations under correctly specified cell-type composition ***π***, and compared the results with methods based on tissue-level data. For iProMix, we considered three hypothesis test strategies: (1) iProMix (*χ*^2^) which uses the theoretical distribution of 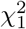 for hypothesis test, (2) iProMix (Permutation) which calculates eFDR using null distribution generated by permuting the label of ACE2, and (3) iProMix (Knockoff) which jointly models the distribution of protein, ACE2 and permuted ACE2. For tissue-level analyses, we considered three additional strategies: (1) regressing ACE2 on other proteins ignoring cell-type composition, (2) regressing ACE2 on other proteins adjusting cell-type composition as a covariate, and (3) regressing main and interactive effects of cell-type composition and ACE2 on other proteins.

In data synthesization, we mimic the real data of CPTAC LUAD to generate 100 tissues. For each tissue *i*, we simulated two major cell types to mimic epithelial and non-epithelial cell components with cell type 1 (epithelial) proportion *π*_*i*_ follows a rescaled Beta distribution as *π*_*i*_ ∼*Beta* (*α=* 3, *β=* 2)/1.4 +0.07. The rescale guarantees cell type 1 is a major cell type with a minimal 7% and mean 50% cell proportion. For ACE2 synthesization, we let the mean level of ACE2 in cell type 1 to be more abundant than in cell type 2, such that *x*_1*i*_ *=* 2 + *e*_1*i*_ and *x*_2*i*_ *=* −2 + *e*_2*i*_, where *e*_2*i*_, *e*_2*i*_ ∼*N* (0, 1). When synthesizing protein data for two cell types (***y***_1*i*_ and ***y***_2*i*_), we let the mean and standard deviation of *P* proteins in cell type 1 and cell type 2 to be randomly generated from *N* (0, 1) and max (0.05, *N* (1, 0.25)). As multiple proteins are simultaneously tested, we further considered the correlation of proteins in data generation, assuming a block-wise correlation structure with 50 proteins in each block. The correlation of two proteins *j* and *l* within the same block follows a autocorrelation structure of *ρ* = 0.6^|*j*−*l*|^. Finally, as the performance of hypothesis tests for cell type 1 may be impacted by the other cell type, we considered five association patterns across the two cell types to comprehensively evaluate the methods:

1. Association in both cell types with the same direction : cor (*x*_1*i*_, *y*_1*i*_) = cor(*x*_2*i*_, *y*_2*i*_) = 0.5,
2. Association in cell type 1 only: cor(*x*_1*i*_, *y*_1*i*_) = 0.5, cor(*x*_2*i*_, *y*_2*i*_) = 0,
3. Association in cell type 2 only: cor(*x*_1*i*_, *y*_1*i*_) = 0, cor((*x*_2*i*_, *y*_2*i*_) = 0.5,
4. Association in both cell types with different directions: cor(*x*_1*i*_, *y*_1*i*_) = 0.5, cor(*x*_2*i*_, *y*_2*i*_) =−0.5, and
5. Association in neither cell type: cor(*x*_1*i*_, *y*_1*i*_) = cor(*x*_2_, *y*_2_) = 0.

We considered a grid of the signal levels to allow the proportion of alternatives ranging 5% to 50% out of a maximum of 3000 simulated proteins. The resulting cell-type specific levels of ACE2 and other proteins cannot be observed, but their weighted averages *x*_*i*_ *= π*_*i*_*x*_1*i*_ + (1 − *π*_*i*_)*x*_2*i*_ for ACE2 and ***y***_*i*_ *= π*_*i*_***y***_1*i*_ +(1 − *π*_*i*_) ***y***_2*i*_ for protein abundance are observed for analysis.

Figure 1 demonstrated the FDR and power of six comparison methods under correctly specified cell-type composition ***π*** across a wide range of signal levels (alternative proportions). When the signals are very abundant that approximately 50% of proteins are associated with ACE2, all methods hold valid FDRs. As the signals become more and more sparse, the FDRs blow up in all tissue-level methods regardless of adjustment for cell-type composition or not. The iProMix (*χ*^2^) performs better than tissue-level methods, but still has inflated FDR when the signal level is <25% of the tests. As the number of proteins associated with ACE2 in real data is believed the be small, iProMix (*χ*^2^) fails to provide valid results. Finally, both iProMix (Permutation) and iProMix (Knockoff) provide valid error control throughout the simulation, and their power are in general comparable.

**Figure 1:**
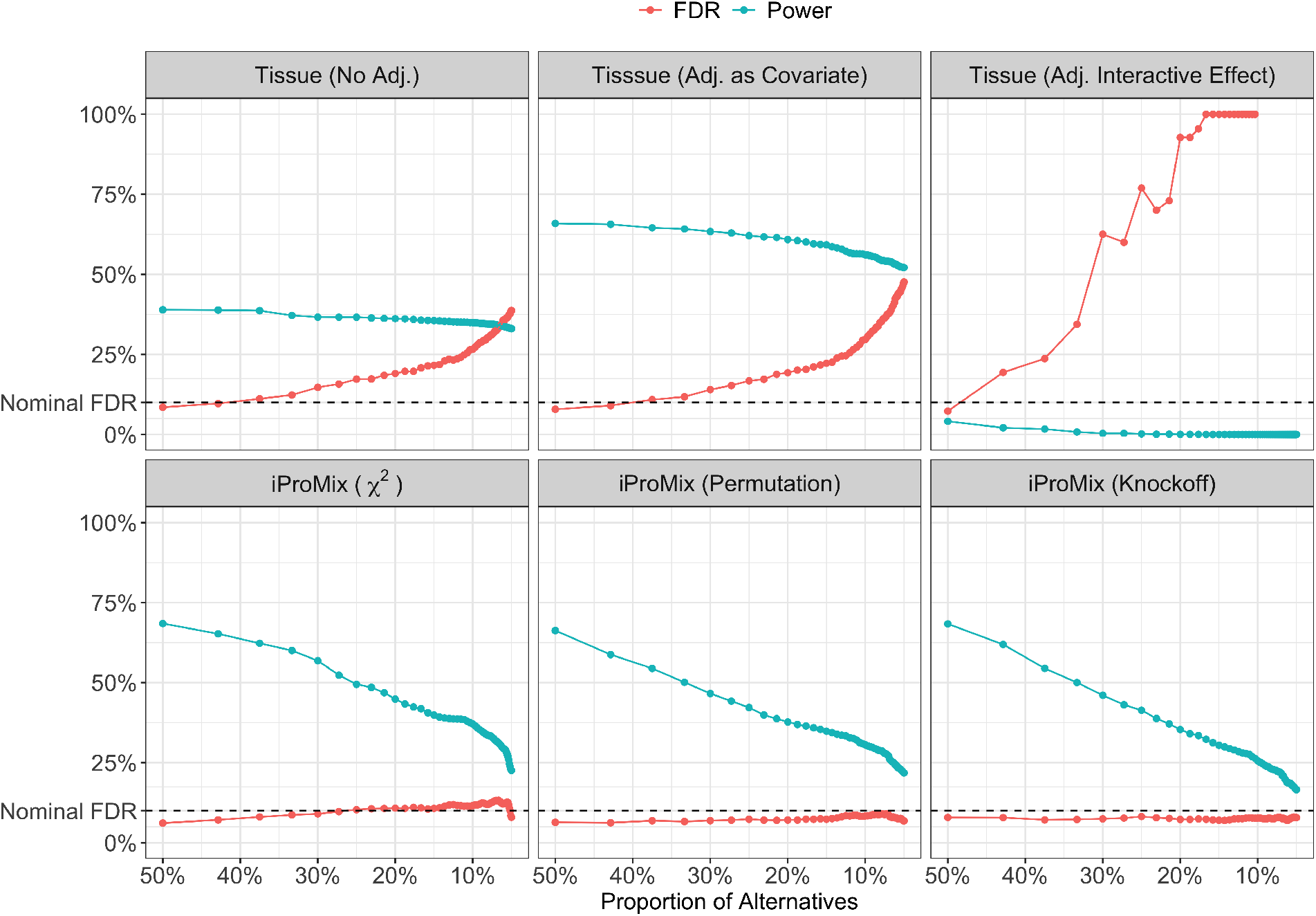
Simulation results for the performance of six comparison methods under correctly specified ***π*** (cell-type composition) across a wide range of signal levels.

### 3.2 Evaluation under misspecification

As in real data cell-type composition is estimated with errors and noises, we further evaluate the performance of iProMix (Permutation) and iProMix (Knockoff) under various scenarios of the misspecified 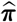. In this simulation, we consider 1000 proteins with five blocks of equal sizes, and each block corresponds to one of the five association patterns mentioned above. We consider various scenarios where 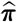 deviates from the true ***π***, such as having scale change, mean shift, reduced correlation, non-linear transformation and including of a third cell type with small cell counts. In details, scale change and mean shift were evaluated by linear transformation of the true ***π***; reduced correlation was achieved by adding random uniformly distributed noises and rescaled to have mean 0.5 and range 0, 1 ; non-linear transformation included the logarithm and exponential transformation of ***π***; and finally, we generated three cell types by splitting cell type 2 into two components with the newly generated cell type 3 taking 10% of the cell counts, and considered the misspecified estimates as a sum of cell type 1 and 3.

The resulting FDR and power of iProMix (Permutation) and iProMix (Knockoff) under misspecified models are listed in Table 1. We found that while both methods have well-controlled FDR rates and comparable power in correctly specified models, iProMix (Permutation) is robust against various model misspecifications but iProMix (Knockoff) is not. iProMix (Knockoff) has inflated FDRs when 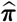 is much greater than true ***π*** (e.g. mean shift from 0.5 to 0.7) and highly impacted power under non-linear transformation and reduced scale. On the contrary, while iProMix (Permutation) is overconservative at true ***π***, it controls FDR and have reasonable power across all mis-specified scenarios under consideration. Therefore, we will applied iProMix (Permutation) to real data where we do not have a perfect knowledge on the cell type composition.

**Table 1:**
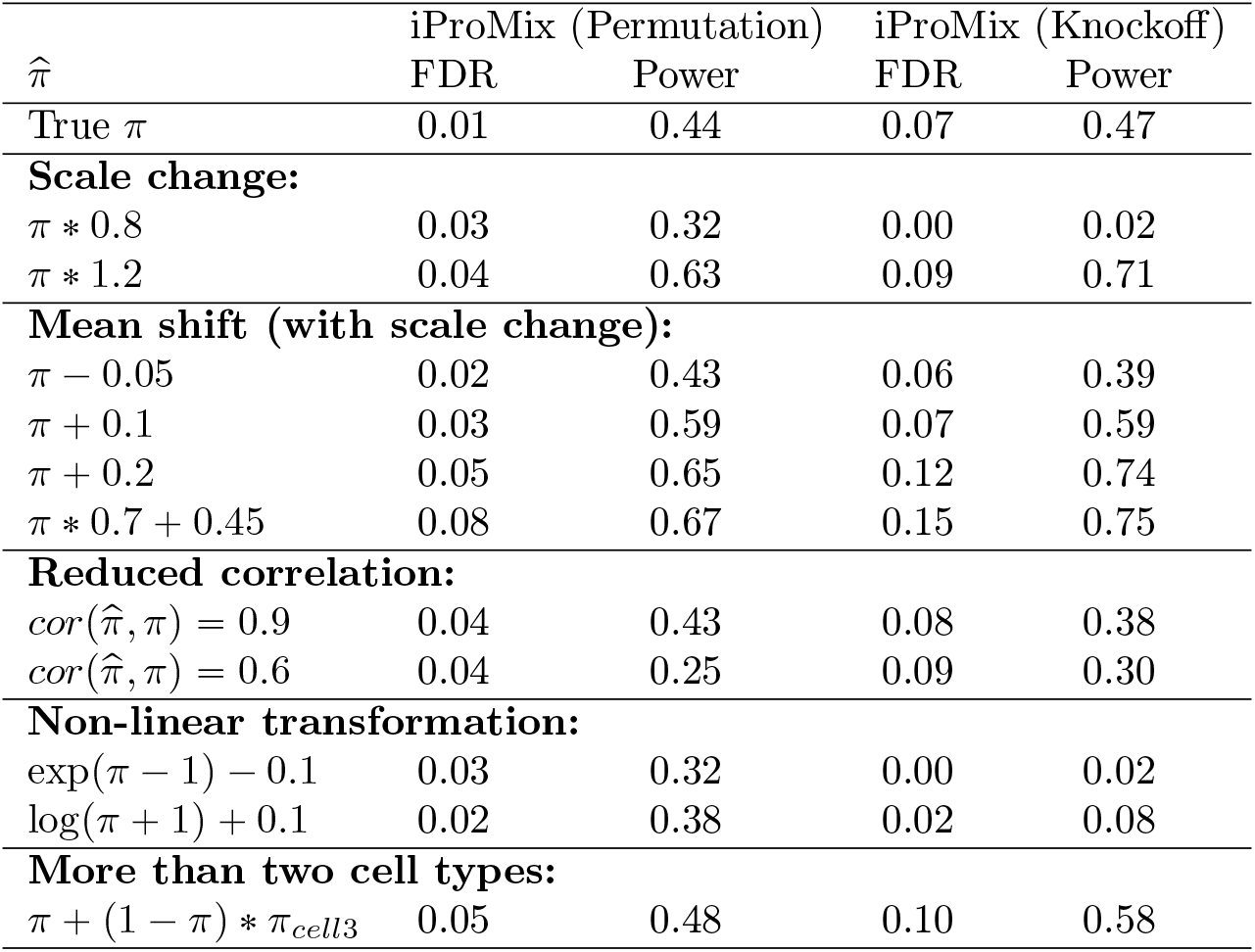
The FDR and power of iProMix (Permutation) and iProMix (Knockoff) methods under mis-specified cell type composition.

## 4 Application to CPTAC LUAD tissue

To identify genes and pathways associated with ACE2 in lung epithelial cells, we applied iProMix to analyze the proteogenomic data sets of 110 normal adjacent tumor (NAT) lung tissue samples from the CPTAC LUAD study [5]. Specifically, the 110 NAT samples were collected from 72 male and 38 female LUAD patients. The transcriptomic and global proteomics data sets contain 18,099 genes and 10,699 proteins respectively. Data prepossessing and quality control were describe in [5]. The gene expression levels and protein abundances of ACE2 in the 110 samples showed no correlation (*p =* 0.254), as illustrated in Figure 2.

**Figure 2:**
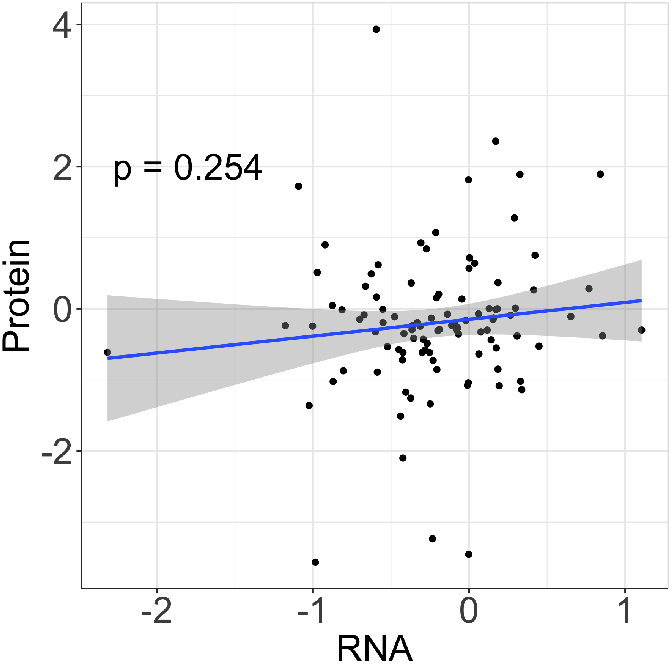
The correlation of the gene expression and protein abundance levels of ACE2 in adjacent normal tissues of lung adenocarcinoma using data from Clinical Proteomic Tumor Analysis Consortium.

### 4.1 Epithelial cell proportion estimation

#### Prior for *π*

First, to obtain a prior estimates of ***π***, we derived epithelial cell scores using xCell [17], a commonly used tool for cell type deconvolution analysis based on RNAseq data. Using cell type specific gene signatures curated from the literature, xCell employs the idea of the single sample gene set enrichment analysis (ssGSEA) scores [32] to capture the relative abundance of each cell type in each sample. The cell type scores from xCell are not direct estimates of cell proportions, and we re-scaled the xCell epithelial scores to have five different mean levels and used re-scaled results as five different sets of priors of ***π*** for iProMix analysis. Specifically, the re-scaling was done by (1) fitting a *Beta* distribution to the xCell epithelial scores; (2) altering the parameter *α* of the *Beta* distribution to generate a new *Beta* distribution with similar shape but different mean, and (3) quantile normalizing the original scores to this new distribution, such that the mean values of the 5 resulting sets of epithelial cell “proportion” vectors evenly spaced from 0.3 to 0.5.

#### Data-driven estimates

For each prior set, we estimated epithelial cell proportions using 126 xCell epithelial signature genes on random bootstrap samples with the iProMix pipeline. We considered a total of 100 bootstrap samples with each being a random drawn of 80% samples, and calculated a truncated mean as our final estimation, which truncated at 5% of tails in both sides and took an average of the remaining 90% estimates. For a total of five priors with mean 0.3-0.5, we have five estimates 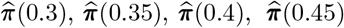 and 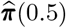 for downstream analysis, and in later models we aggregate identifications with all priors.

The performance of estimated epithelial cell proportions is presented in Figure 3, and compared to xCell scores (prior information). The correlations of 126 epithelial signature genes with estimated epithelial proportions are higher than the correlation with xCell scores, with a median correlation improved from 0.246 in xCell to 0.292 − 0.296 in iProMix. A total of 82% epithelial signatures were positively correlated with the xCell scores, while all epithelial signatures were positively correlated with the iProMix estimates. This indicated the 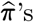 were more relevant to epithelial cells than xCell scores. In Figure 3(b), the correlations among iProMix estimates and with xCell scores were contrasted. A high correlation among iProMix estimates were observed (r = 0.995-1; p-value < 2e-16), indicating a highly consistent estimation with different priors. Their correlation with xCell score is low (r= 0.217-0.226; p-value = 0.023 -0.030).

**Figure 3:**
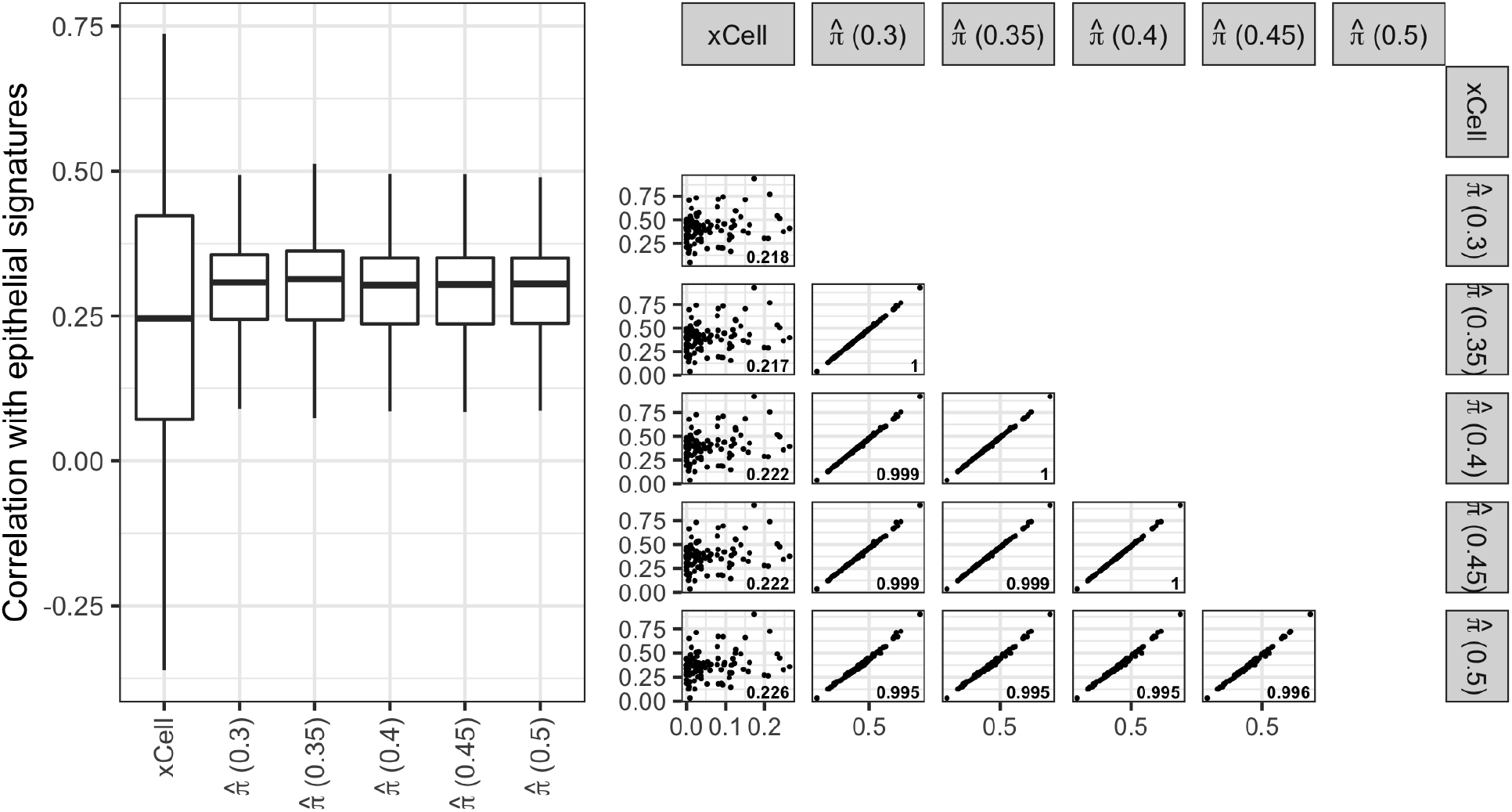
Evaluation of epithelial cell proportion estimates for CPTAC NAT lung tissue samples. (a). Boxplot of the correlation between xCell epithelial score, estimated epithelial cell proportions from the iProMix pipeline using different prior parameters, and the expression levels of 126 individual epithelial cell signature genes from xCell [17]. (b). Correlation between xCell epithelial score and estimated epithelial cell proportions from the iProMix pipeline.

### 4.2 Genes identification

With estimated epithelial cell proportions, we applied iProMix to the proteomics data of CPTAC LUAD adjacent normal tissues to identify epithelial-specific associations. We first considered the protein abundance of ACE2 with each of the 10,698 remaining proteins quantified in the data. The analysis included age and smoking status as covariates, and stratified by sex to allow sex-specific effects as ACE2 is a gene located on X chromosome. Then, we repeated the same analysis for the expression level of ACE2 with all 10,699 proteins in order to further understand these genes for their signals at expression levels.

We employed iProMix (Permutation) procedure to estimate the cell-type specific correlations and identify proteins at a 10% eFDR for each of the five 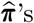. We then considered the consensus voting across the five 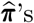 to declare significant proteins that were significant in at least 50% of the tests. For comparison purposes, we also modeled the association between ACE2 and other proteins at tissue level with three strategies (1) regression with no adjustment of cell type composition, (2) regression adjusting cell type composition as a covariate, and regressing the interactive effect of ACE2 and cell type composition on other proteins.

Table 2 presented the number of genes identified by each approach, as well as names of significant genes. While no genes were identified to be associated with ACE2 protein levels at tissue-level models, iProMix identified RNASE1 in men and 20 genes in women. However, none of the identified genes were replicated at the expression level of ACE2 at FDR 10%. The expression level of ACE2 were only identified to be associated with the interactive effect of JADE1 and epithelial cell proportion in men.

**Table 2:** Number of genes identified at FDR 0.1 using tissue-level methods and iProMix, as well as name of identified genes. * The 20 genes include PBXIP1, SGPL1, FAM50B, OTUB1, TMCC1, DDX11, RETREG3, HACD3, HPS1, RNF167, PROX1, IKZF2, SOX4, FAM3A, SRD5A3, CFAP157, METAP1D, FGD1, SPOCD1 and NDUFAF8.

| | Tissue (No Adj.) | Tissue (Adj. as Cov) | Tissue ($\pi \times$ ACE2) | iProMix (Epithelial) |
| --- | --- | --- | --- | --- |
| **ACE2 Protein with all other proteins:** |  |  |  |  |
| Male | 0 | 0 | 0 | 1 (RNASE1) |
| Female | 0 | 0 | 0 | 20* |
| **ACE2 RNA with all proteins:** |  |  |  |  |
| Male | 0 | 0 | 1 (JADE1) | 0 |
| Female | 0 | 0 | 0 | 0 |

### 4.3 Pathway enrichment for LUAD proteomics

We considered 50 pathways from Hallmark database [31] including IFN*α* and IFN*γ* to identify pathways associated with ACE2 in epithelial cell component. iProMix was applied with each of the five pre-estimated 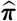, and the resulting p-values were aggregated using Cauchy combination [33] to get the final p-values across all 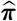. The BY procedure was then applied to the combined p-values to identify pathways associated with ACE2 in epithelial cells. For comparison purposes, we also performed tissue-level analysis and compare for BY controlled pathway discoveries, and if 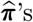 were used, the same Cauchy combination was applied to combine p-values for comparison.

The most significant pathways showing positive association with ACE2 protein abundances in epithelial cells were IFN*α*/*γ* pathways (Figure S1). These associations were also observed by looking at the expression levels of ACE2 (Figure S2). Figure 4 compared the association p-values of iProMix with tissue-level methods for IFN*γ* and INF*α* pathways. When the cell type composition was not adjusted, in both men and women, at both gene expression and protein abundance levels, the IFN*α* response pathway was not identified. After adjusting for estimated epithelial cell composition as a covariate, the IFN*α* response was positively associated with ACE2 protein abundance level in men. The interaction model identified the effects of ACE2 on IFN*α* response pathway differ by epithelial cell composition in men at its protein level and in women at both RNA and protein levels. Finally, iProMix identified that both ACE2 protein and expression levels in epithelial cell component were positively associated with IFN*α* response pathway in men, and both were negatively associated with IFN*α* response pathway in women. The results for the IFN*γ* response pathway was very similar to INF*α* that we were able to identify positive associations between ACE2 and INF*γ* in men and negative associations in women at both protein and RNA levels.

**Figure 4:**
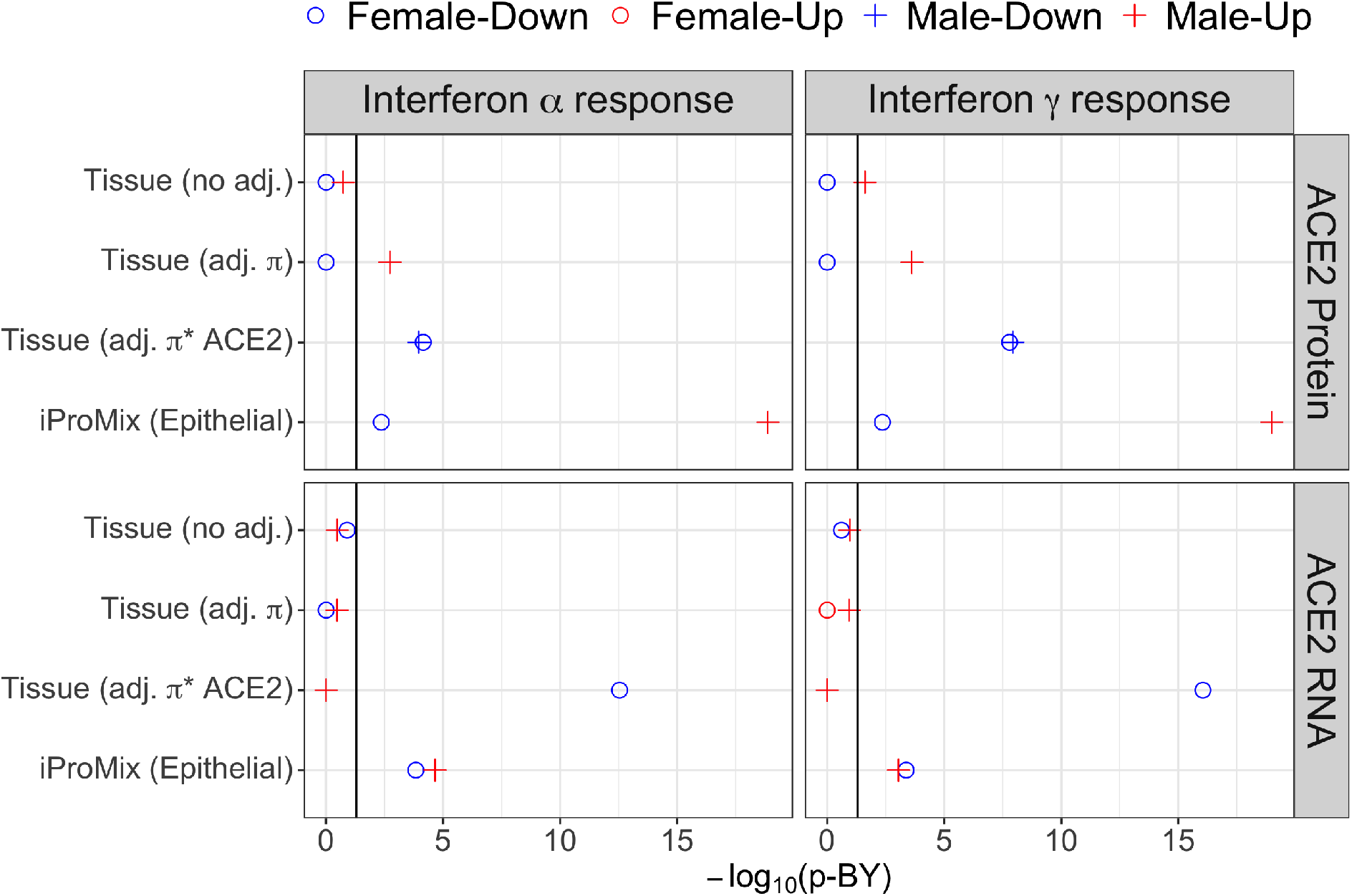
Pathway enrichment results for IFN*γ* and INF*α* pathways using tissue-level models and iProMix for men and women. Sex differences in immune responses was observed. Both IFN*γ* and INF*α* pathways were significantly positively associated with both ACE2 expression and protein levels in epithelial cell component in men. These two pathways were negatively associated with ACE2 expression and protein levels in epithelial cell component in women.

More intriguingly, in the iProMix results, the positive associations between ACE2 and IFN-*α*/*γ* were observed only in the tissue samples from male patients, while the association direction is negative in samples from female patients. The sex difference for the association between *ACE*2 and IFN pathways was uniquely identified in our study and may provide potential explanations for the sex difference in immune response to SARS-Cov-2. To provider further understanding of the key players that contribute to sex difference, we further contrasted men and women for the rank of signed LRT scores of the genes within these two pathways. The one with the most striking sex difference was TRIM14 (Figure 5).

**Figure 5:**
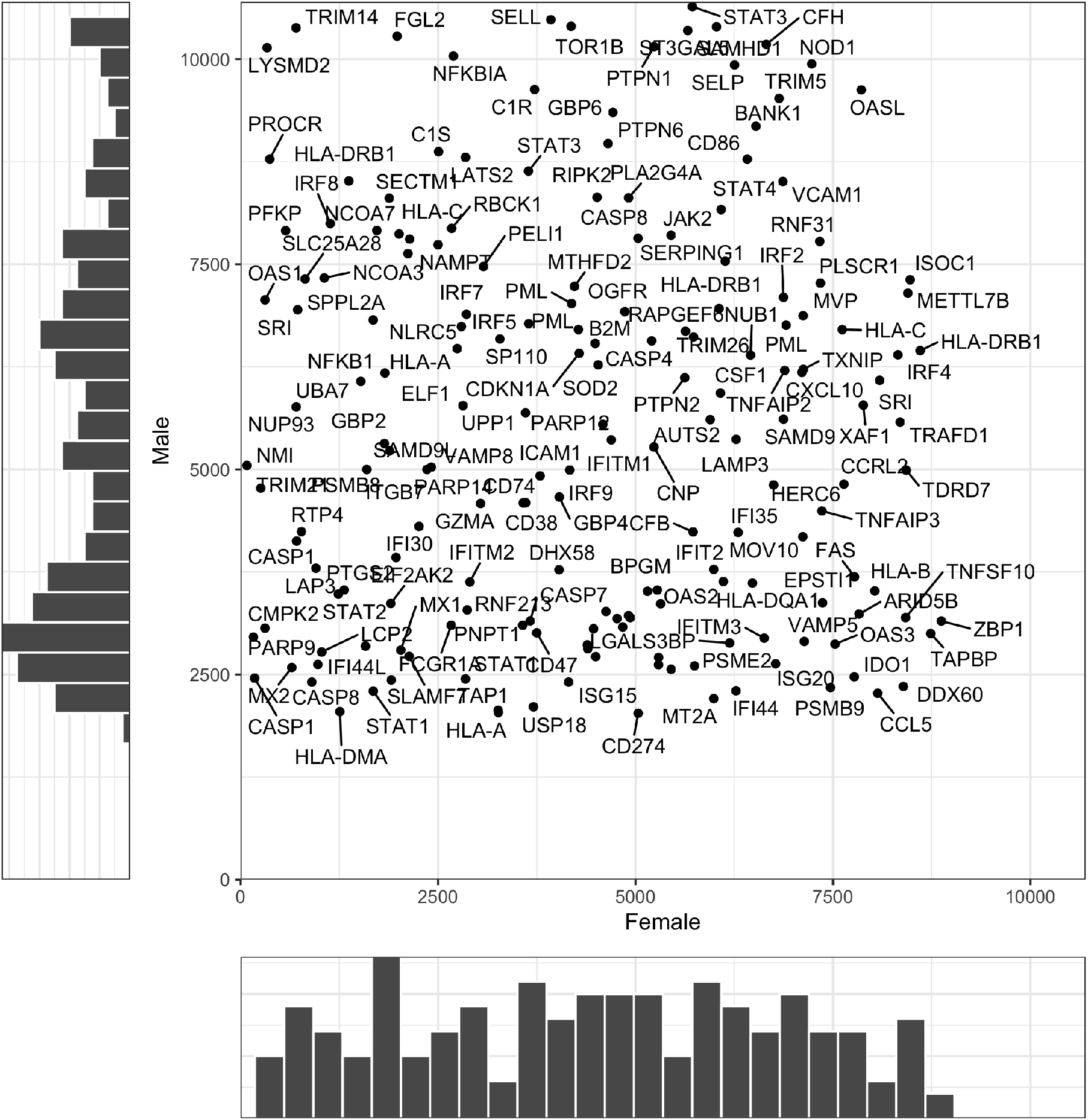
A contrast of ranks of signed LRT scores for genes in INF*α* and INF*γ* pathways in male and female patients. The upper left corner (TRIM14) presents that genes that contribute the most sex differences in their associations with ACE2 protein levels.

## 5 Discussion

In this work, we develop a new method *iProMix* to test for cell-type specific associations among genes/proteins based on bulk tissue proteogenomic profiles. Besides providing association inferences, iProMix is able to take (noisy) cell type percentage estimates as priors and deliver improved percentage estimates in the final results. In the simulation study, we show that *iProMix* properly controls the FDRs and is more powerful than tissue-level analysis for detecting the cell-type specific associations. When iProMix is applied to the CPTAC LUAD proteogenomic data to study ACE2 related genes/proteins, it not only validates the associations between ACE2 and IFN pathways in epithelial cells of human lung tissues, but also revealed novel sex-specific effects.

ACE2 has been suggested to be an IFN stimulated gene in human airway epithelial cells by a recent study based on single cell RNAseq data and *in vitro* experiments [4]. Thus, the fact that iProMix detected IFN*α* and INF*γ* as the most significant pathways associated with ACE2 suggests the proposed method is able to capture real biological signals in the bulk tissue data. Moreover, intriguingly, in the iProMix results, the positive associations between ACE2 and IFN*α* and INF*γ* were observed only in male patients, while the association was negative in female patients. It has been reported that COVID-19 produced more severe symptoms and higher mortality among men than women [34, 35]. Studies examining the antibody titres and plasma cytokines in COVID-19 patients revealed a sex difference in immune response to COVID-19 virus [36]. The result in our analysis further suggested that the sex difference in immune response to COVID-19 could be partly due to the different regulation patterns between IFN and ACE2 in lung epithelial cells in males and females. For genes in the INF*α* and INF*γ* pathways, TRIM14 was identified to show the most striking sex difference. TRIM14 is a mitochondrial adaptor that facilitates retinoic acid-inducible gene-I–like receptor-mediated innate immune response [37]. It has been reported as a key regulator of the Type I IFN response in other virus infections [38]. Therefore, by applying the new tool iProMix to the real data, we are able to detect promising biological signals in the proteomics that are otherwise missed by using tissue level methods. Nevertheless, cautions should be given in scientific interpretation the results. Analyses were performed on adjacent normal tissue of lung cancer, which may have different protein levels and associations from normal tissues of non-cancer patients. Also, A comparison of men and women should be aware that they have different sample sizes in our data which may affect the power and accuracy in detecting their signals.

The method is applicable to a wide range of studies where the interest is on the dependence of two genes in single or multiple -omic data types that are impacted by cell type composition. For example, analysis on the interrelationship of genes within or across epigenomics, transcriptomics, proteomics and metabolisms can employ iProMix framework for cell-type specific associations. Extension of the tool to more than two cell types is theoretical feasible but the analytical performance for rare cell types needs careful evaluation. Joint consideration of multiple genes, such as via Gaussian graphic models, for their cell-type specific conditional associations is another direction for extension of the method, in which regularization should be further investigated for a good performance. Finally, gene identification is currently achieved by consensus voting of results from multiple priors, and an exploration of more strategies for aggregating results may further improve the study power.

Software implementing the proposed iProMix is available on R CRAN at https://cran.r-project.org/iProMix [To be announced upon acceptance], the data containing genes and pathways identified by iProMix is available on Github at [To be announced upon acceptance].

## Acknowledgements

This work has been supported by National Institute of Health grant 5U24CA210993 to XS, JJ and PW; by National Institute of Health grant P30CA196521 to XS and JJ.

## Supplementary Materials

**Figure S1:**
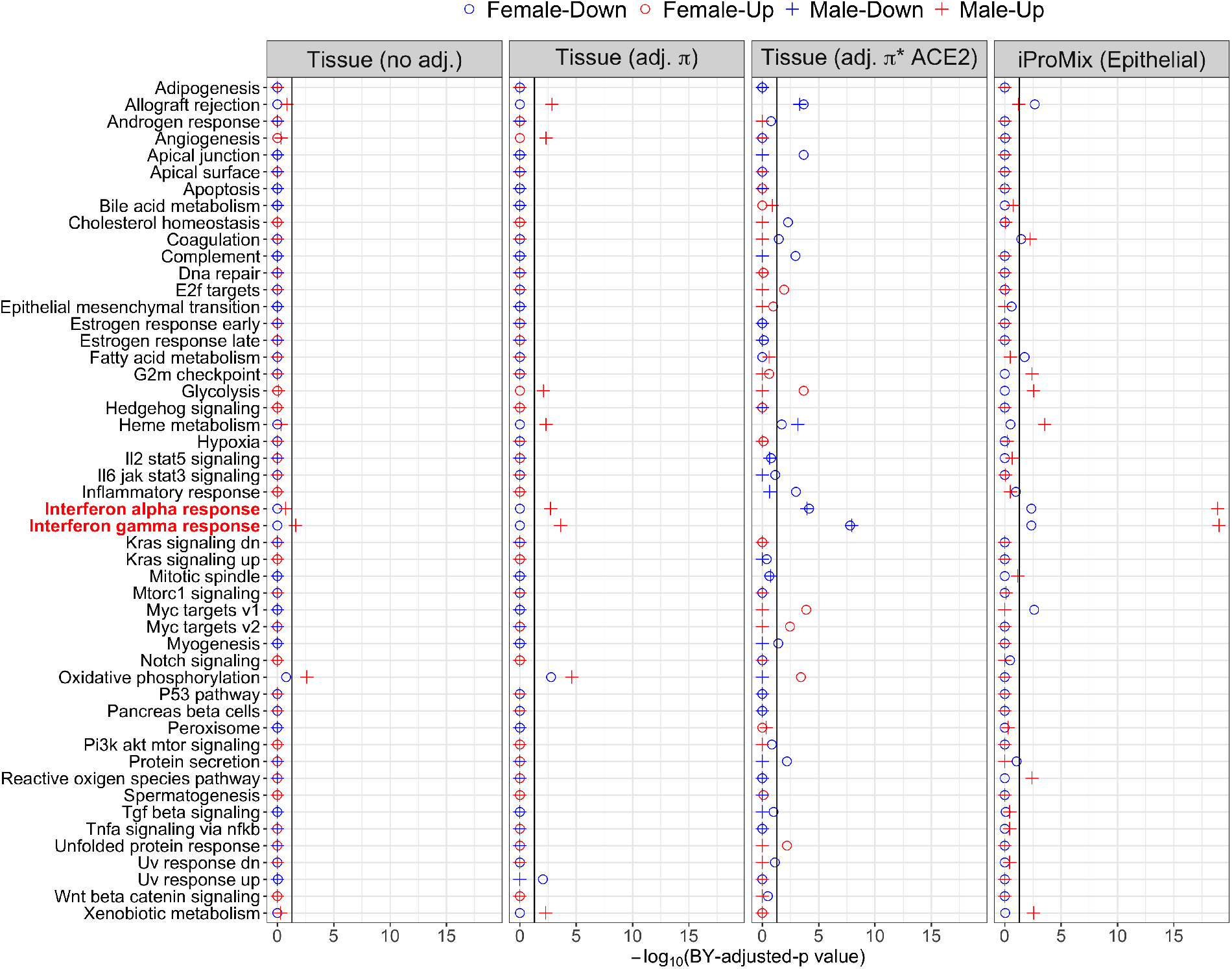
Pathway enrichment analysis for 50 Hallmark pathways associated with ACE2 protein level.

**Figure S2:**
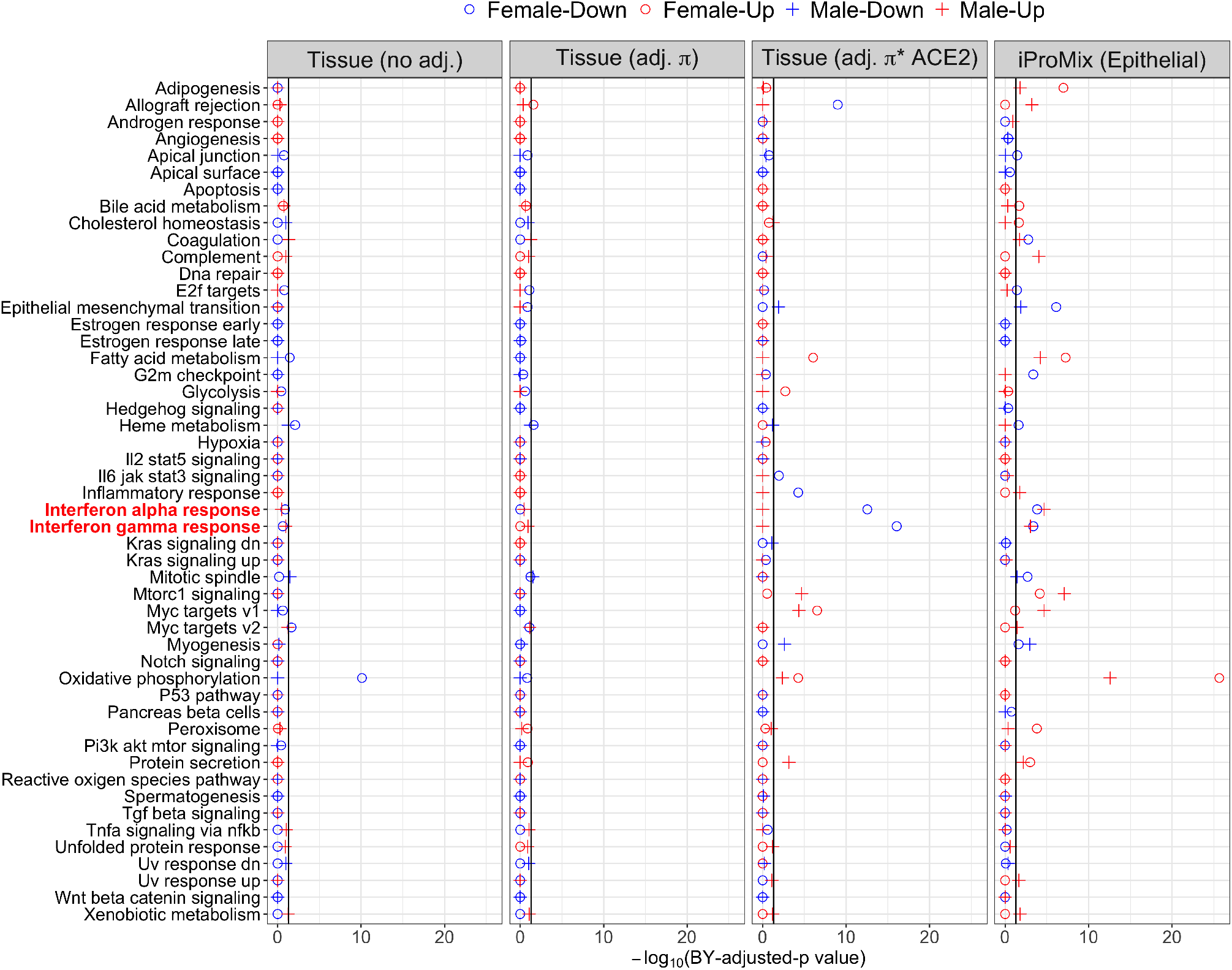
Pathway enrichment analysis for 50 Hallmark pathways associated with ACE2 RNA level.

### Identifiability of Θ

Suppose we consider an alternative sets of parameter 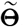, and then the iProMix model is identifiable if 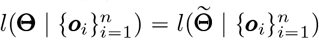 for any observed data ***o***_*i*_’s implies 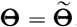. Suppose the 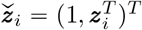 includes the intercept and *P* dimensional covariates. As the normal distribution is identifiable, the mean function of the ACE2 can be written as

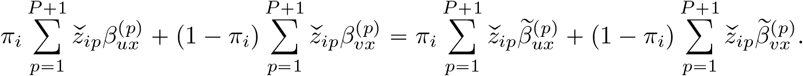

This equation can be rewritten into a matrix form as follows:

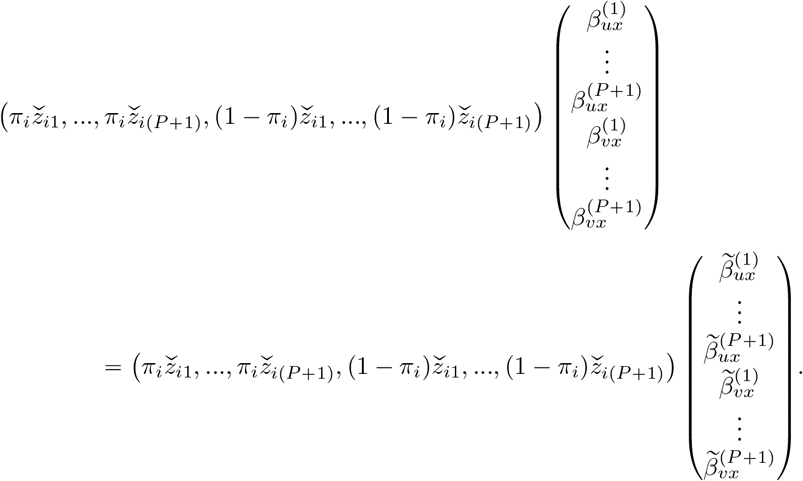

By combining these *n* equations, the first component 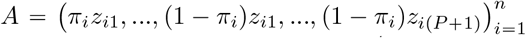 is a *n* × 2(*P* + 1) matrix with a rank of 2(*P* + 1) (full column rank). *A* has a left inverse *A* ^−1^, and multiplying by *A* ^−1^ from the left on both sides, we obtain 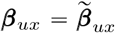 and 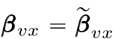. Similarly, we were able to prove 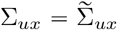 and 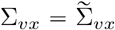, and the rest of the paramters for the protein. As a result, 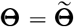 and iProMix is identifiabile.

## Notes

### Competing Interest Statement

The authors have declared no competing interest.

## References

[1] Markus Hoffmann, Hannah Kleine-Weber, Simon Schroeder, Nadine Krüger, Tanja Herrler, Sandra Erichsen, Tobias S Schiergens, Georg Herrler, Nai-Huei Wu, Andreas Nitsche, et al. Sars-cov-2 cell entry depends on ace2 and tmprss2 and is blocked by a clinically proven protease inhibitor. Cell, 2020.

[2] Waradon Sungnak, Ni Huang, Christophe Bécavin, Marijn Berg, Rachel Queen, Monika Litvinukova, Carlos Talavera-López, Henrike Maatz, Daniel Reichart, Fotios Sampaziotis, et al. Sars-cov-2 entry factors are highly expressed in nasal epithelial cells together with innate immune genes. Nature medicine, 26(5):681–687, 2020.

[3] Keiji Kuba, Yumiko Imai, Shuan Rao, Hong Gao, Feng Guo, Bin Guan, Yi Huan, Peng Yang, Yanli Zhang, Wei Deng, et al. A crucial role of angiotensin converting enzyme 2 (ace2) in sars coronavirus– induced lung injury. Nature medicine, 11(8):875–879, 2005.

[4] Carly GK Ziegler, Samuel J Allon, Sarah K Nyquist, Ian M Mbano, Vincent N Miao, Constantine N Tzouanas, Yuming Cao, Ashraf S Yousif, Julia Bals, Blake M Hauser, et al. Sars-cov-2 receptor ace2 is an interferon-stimulated gene in human airway epithelial cells and is detected in specific cell subsets across tissues. Cell, 2020.

[5] Michael A Gillette, Shankha Satpathy, Song Cao, Saravana M Dhanasekaran, Suhas V Vasaikar, Karsten Krug, Francesca Petralia, Yize Li, Wen-Wei Liang, Boris Reva, et al. Proteogenomic characterization reveals therapeutic vulnerabilities in lung adenocarcinoma. Cell, 182(1):200–225, 2020.

[6] David J Clark, Saravana M Dhanasekaran, Francesca Petralia, Jianbo Pan, Xiaoyu Song, Yingwei Hu, Felipe da Veiga Leprevost, Boris Reva, Tung-Shing M Lih, Hui-Yin Chang, et al. Integrated proteogenomic characterization of clear cell renal cell carcinoma. Cell, 179(4):964–983, 2019.

[7] Francesca Petralia, Nicole Tignor, Boris Reva, Pichai Raman, Shrabanti Chowdhury, Dmitry Rykunov, Azra Krek, Weiping Ma, Jiayi Ji, Xiaoyu Song, et al. Integrated proteogenomic characterization across seven histological types of pediatric brain tumors, 2020.

[8] Xiaoping Su, Li Zhang, Jianping Zhang, Funda Meric-Bernstam, and John N Weinstein. Purityest: estimating purity of human tumor samples using next-generation sequencing data. Bioinformatics, 28(17):2265–2266, 2012.

[9] David Venet, F Pecasse, Carine Maenhaut, and Hugues Bersini. Separation of samples into their constituents using gene expression data. Bioinformatics, 17(suppl_1):S279–S287, 2001.

[10] Timo Erkkilä, Saara Lehmusvaara, Pekka Ruusuvuori, Tapio Visakorpi, Ilya Shmulevich, and Harri Lähdesmäki. Probabilistic analysis of gene expression measurements from heterogeneous tissues. Bioinformatics, 26(20):2571–2577, 2010.

[11] Christopher R Bolen, Mohamed Uduman, and Steven H Kleinstein. Cell subset prediction for blood genomic studies. BMC bioinformatics, 12(1):258, 2011.

[12] Shai S Shen-Orr, Robert Tibshirani, Purvesh Khatri, Dale L Bodian, Frank Staedtler, Nicholas M Perry, Trevor Hastie, Minnie M Sarwal, Mark M Davis, and Atul J Butte. Cell type–specific gene expression differences in complex tissues. Nature methods, 7(4):287, 2010.

[13] Jason E Shoemaker, Tiago JS Lopes, Samik Ghosh, Yukiko Matsuoka, Yoshihiro Kawaoka, and Hiroaki Kitano. Cten: a web-based platform for identifying enriched cell types from heterogeneous microarray data. BMC genomics, 13(1):460, 2012.

[14] Kosuke Yoshihara, Maria Shahmoradgoli, Emmanuel Martínez, Rahulsimham Vegesna, Hoon Kim, Wandaliz Torres-Garcia, Victor Treviño, Hui Shen, Peter W Laird, Douglas A Levine, et al. Inferring tumour purity and stromal and immune cell admixture from expression data. Nature communications, 4:2612, 2013.

[15] Scott L Carter, Kristian Cibulskis, Elena Helman, Aaron McKenna, Hui Shen, Travis Zack, Peter W Laird, Robert C Onofrio, Wendy Winckler, Barbara A Weir, et al. Absolute quantification of somatic dna alterations in human cancer. Nature biotechnology, 30(5):413, 2012.

[16] Aaron M Newman, Chih Long Liu, Michael R Green, Andrew J Gentles, Weiguo Feng, Yue Xu, Chuong D Hoang, Maximilian Diehn, and Ash A Alizadeh. Robust enumeration of cell subsets from tissue expression profiles. Nature methods, 12(5):453–457, 2015.

[17] Dvir Aran, Zicheng Hu, and Atul J Butte. xcell: digitally portraying the tissue cellular heterogeneity landscape. Genome biology, 18(1):220, 2017.

[18] Elior Rahmani, Regev Schweiger, Liat Shenhav, Theodora Wingert, Ira Hofer, Eilon Gabel, Eleazar Eskin, and Eran Halperin. Bayescce: a bayesian framework for estimating cell-type composition from dna methylation without the need for methylation reference. Genome biology, 19(1):1–18, 2018.

[19] Ankur Chakravarthy, Andrew Furness, Kroopa Joshi, Ehsan Ghorani, Kirsty Ford, Matthew J Ward, Emma V King, Matt Lechner, Teresa Marafioti, Sergio A Quezada, et al. Pan-cancer deconvolution of tumour composition using dna methylation. Nature communications, 9(1):1–13, 2018.

[20] Zeya Wang, Shaolong Cao, Jeffrey S Morris, Jaeil Ahn, Rongjie Liu, Svitlana Tyekucheva, Fan Gao, Bo Li, Wei Lu, Ximing Tang, et al. Transcriptome deconvolution of heterogeneous tumor samples with immune infiltration. iScience, 9:451–460, 2018.

[21] Xuran Wang, Jihwan Park, Katalin Susztak, Nancy R Zhang, and Mingyao Li. Bulk tissue cell type deconvolution with multi-subject single-cell expression reference. Nature communications, 10(1):1–9, 2019.

[22] Gregory J Hunt, Saskia Freytag, Melanie Bahlo, and Johann A Gagnon-Bartsch. dtangle: accurate and robust cell type deconvolution. Bioinformatics, 35(12):2093–2099, 2019.

[23] Aaron M Newman, Chloé B Steen, Chih Long Liu, Andrew J Gentles, Aadel A Chaudhuri, Florian Scherer, Michael S Khodadoust, Mohammad S Esfahani, Bogdan A Luca, David Steiner, et al. Determining cell type abundance and expression from bulk tissues with digital cytometry. Nature biotechnology, 37(7):773–782, 2019.

[24] Francesca Petralia, Li Wang, Jie Peng, Arthur Yan, Jun Zhu, and Pei Wang. A new method for constructing tumor specific gene co-expression networks based on samples with tumor purity heterogeneity. Bioinformatics, 34(13):i528–i536, 2018.

[25] Jaeil Ahn, Ying Yuan, Giovanni Parmigiani, Milind B Suraokar, Lixia Diao, Ignacio I Wistuba, and Wenyi Wang. Demix: deconvolution for mixed cancer transcriptomes using raw measured data. Bioinformatics, 29(15):1865–1871, 2013.

[26] Margaret KR Donovan, Agnieszka D’Antonio-Chronowska, Matteo D’Antonio, and Kelly A Frazer. Cellular deconvolution of gtex tissues powers discovery of disease and cell-type associated regulatory variants. Nature communications, 11(1):1–14, 2020.

[27] Xiangyu Luo, Can Yang, and Yingying Wei. Detection of cell-type-specific risk-cpg sites in epigenome-wide association studies. Nature communications, 10(1):1–12, 2019.

[28] Xiao-Li Meng and Donald B Rubin. Maximum likelihood estimation via the ecm algorithm: A general framework. Biometrika, 80(2):267–278, 1993.

[29] John D Storey. A direct approach to false discovery rates. Journal of the Royal Statistical Society: Series B (Statistical Methodology), 64(3):479–498, 2002.

[30] Matthew E Ritchie, Belinda Phipson, D. Wu, Yifang Hu, Charity W Law, Wei Shi, and Gordon K Smyth. limma powers differential expression analyses for rna-sequencing and microarray studies. Nucleic acids research, 43(7):e47–e47, 2015.

[31] Arthur Liberzon, Chet Birger, Helga Thorvaldsdóttir, Mahmoud Ghandi, Jill P Mesirov, and Pablo Tamayo. The molecular signatures database hallmark gene set collection. Cell systems, 1(6):417–425, 2015.

[32] David A Barbie, Pablo Tamayo, Jesse S Boehm, So Young Kim, Susan E Moody, Ian F Dunn, Anna C Schinzel, Peter Sandy, Etienne Meylan, Claudia Scholl, et al. Systematic rna interference reveals that oncogenic kras-driven cancers require tbk1. Nature, 462(7269):108–112, 2009.

[33] Yaowu Liu and Jun Xie. Cauchy combination test: a powerful test with analytic p-value calculation under arbitrary dependency structures. Journal of the American Statistical Association, 115(529):393– 402, 2020.

[34] Nanshan Chen, Min Zhou, Xuan Dong, Jieming Qu, Fengyun Gong, Yang Han, Yang Qiu, Jingli Wang, Ying Liu, Yuan Wei, et al. Epidemiological and clinical characteristics of 99 cases of 2019 novel coronavirus pneumonia in wuhan, china: a descriptive study. The lancet, 395(10223):507–513, 2020.

[35] Catherine Gebhard, Vera Regitz-Zagrosek, Hannelore K Neuhauser, Rosemary Morgan, and Sabra L Klein. Impact of sex and gender on covid-19 outcomes in europe. Biology of sex differences, 11:1–13, 2020.

[36] Takehiro Takahashi, Mallory K Ellingson, Patrick Wong, Benjamin Israelow, Carolina Lucas, Jon Klein, Julio Silva, Tianyang Mao, Ji Eun Oh, Maria Tokuyama, et al. Sex differences in immune responses that underlie covid-19 disease outcomes. Nature, 588(7837):315–320, 2020.

[37] Zhuo Zhou, Xue Jia, Qinghua Xue, Zhixun Dou, Yijie Ma, Zhendong Zhao, Zhengfan Jiang, Bin He, Qi Jin, and Jianwei Wang. Trim14 is a mitochondrial adaptor that facilitates retinoic acid-inducible gene-i–like receptor-mediated innate immune response. Proceedings of the National Academy of Sciences, 111(2):E245–E254, 2014.

[38] Caitlyn T Hoffpauir, Samantha L Bell, Kelsi O West, Tao Jing, Allison R Wagner, Sylvia Torres-Odio, Jeffery S Cox, A Phillip West, Pingwei Li, Kristin L Patrick, et al. Trim14 is a key regulator of the type i ifn response during mycobacterium tuberculosis infection. The Journal of Immunology, 205(1):153–167, 2020.

